# Light Guiding in Scattering Media Using Focused Laser-Induced Acoustic Waves with an Open Elliptical Reflector

**DOI:** 10.64898/2026.09.17.752288

**Authors:** Yijie Liu, Keitaro Shimada, Kenta Kitamura, Santiago Barrera, Maosen Ye, Ayumu Ishijima, Keiichi Nakagawa

**Affiliations:** Department of Bioengineering, The University of Tokyo, Tokyo 113-8656, Japan; Department of Precision Engineering, The University of Tokyo, Tokyo 113-8656, Japan; Department of Engineering, Friedrich-Alexander-Universität Erlangen-Nürnberg, 91054 Erlangen, Germany

## Abstract

Light scattering in biological tissue limits deep optical imaging. Acoustic waves can modulate optical properties and enhance photon delivery, but source-positioning requirements can complicate integration with other imaging modalities. Here, we present an open elliptical reflector that focuses laser-induced acoustic waves at a spatially separated target while maintaining open optical access. We characterized the spatiotemporal optical modulation induced by the localized acoustic focus. The system increased the local transmitted-light intensity by a factor of 2.57-fold through a 2-mm-thick 1% intralipid hydrogel and enhanced fluorescence signal under two-photon excitation through a 1.5-mm-thick scattering sample. These results demonstrate spatially separated, reflector-based acoustically assisted light guiding for enhanced light delivery and fluorescence excitation through scattering media.

---

Light propagation in biological tissue is strongly affected by scattering, which attenuates and spatially redistributes excitation photons and thereby limits high-resolution optical imaging at increasing tissue depths. This limitation is particularly critical in neuronal imaging, where high-resolution visualization of brain microstructure and morphology requires efficient light delivery through scattering tissue. Among optical techniques, two-photon fluorescence microscopy (TPM), typically using near-infrared excitation in the 700–1100 nm range, provides high spatial resolution and molecular specificity with deeper penetration than one-photon microscopy. Nevertheless, scattering progressively reduces the number of excitation photons reaching the focus and increases the background generated outside the focal volume, fundamentally limiting the achievable imaging depth to several hundred micrometers in typical in vivo brain imaging and to approximately 1 mm under optimized conditions [1].

To overcome this limitation, invasive optical delivery devices such as gradient-index lenses and optical fibers have been used to access deep brain regions in animal studies [2-4]. Despite their effectiveness, these approaches require physical insertion into tissue and may cause tissue damage or perturbation. As a noninvasive alternative, wave-modulation techniques, including wavefront shaping with spatial light modulators [5,6], can in principle compensate for scattering by tailoring the incident wavefront. In contrast to wavefront-shaping methods that rely on characterization and active compensation of scattering-induced optical distortions, acoustically assisted light guiding provides a complementary approach by transiently modifying the local light-propagation environment without explicit wavefront reconstruction [7-10]. In these approaches, acoustic waves or pressure fields generated by piezoelectric ultrasound transducers or laser sources induce localized modulation of the optical properties within the medium. These modulated regions can guide and redirect light over millimeter-scale distances, thereby enhancing optical delivery without the physical insertion of optical probes or iterative wavefront optimization.

However, some previously reported acoustically assisted light guiding configurations impose strict positional constraints between the acoustic source and the target region. This requirement limits the available optical-access geometry and introduces potential spatial interference when integrating these methods with other imaging modalities, such as two-photon microscopy. Further flexibility in separating the modulation region from the acoustic source would facilitate integration with real-time multimodal imaging systems. A previously reported approach used a parabolic reflector to shape transversely propagating ultrasound into an extended column-shaped field for transient light guiding, but required acoustic components on both sides of the sample, thereby limiting optical access and system integration [9].

In this Letter, we introduce a finite-thickness partial elliptical acoustic reflector that can redirect laser-induced acoustic waves from one focal region to another, producing localized acoustic focusing at a selected position within a scattering medium [11]. The reflector design is motivated by three-dimensional ellipsoidal acoustic reflectors [12,13], in which waves originating near one focus are redirected by the curved surface toward the other focus. This configuration spatially separates acoustic-wave generation from the target region while maintaining open optical access. We experimentally characterized the spatiotemporal acoustic focusing and demonstrated enhanced light transmission and fluorescence excitation through scattering media.

In practice, the laser-induced acoustic source and the redirected acoustic focus occupy finite spatial regions rather than ideal points; we therefore refer to F1 and F2 as the first and second focal regions, respectively. Acoustic waves generated near F1 are reflected by the elliptical surface and converge near F2 (Fig. 1(a)). Because the reflector has a finite thickness and an elliptical cross-section, acoustic focusing is governed primarily by the in-plane elliptical geometry, while confinement in the out-of-plane direction is limited. This design therefore produces a localized focal volume rather than an ideal point focus, while maintaining an open geometry for optical access and integration with the imaging system.

**Fig. 1.**
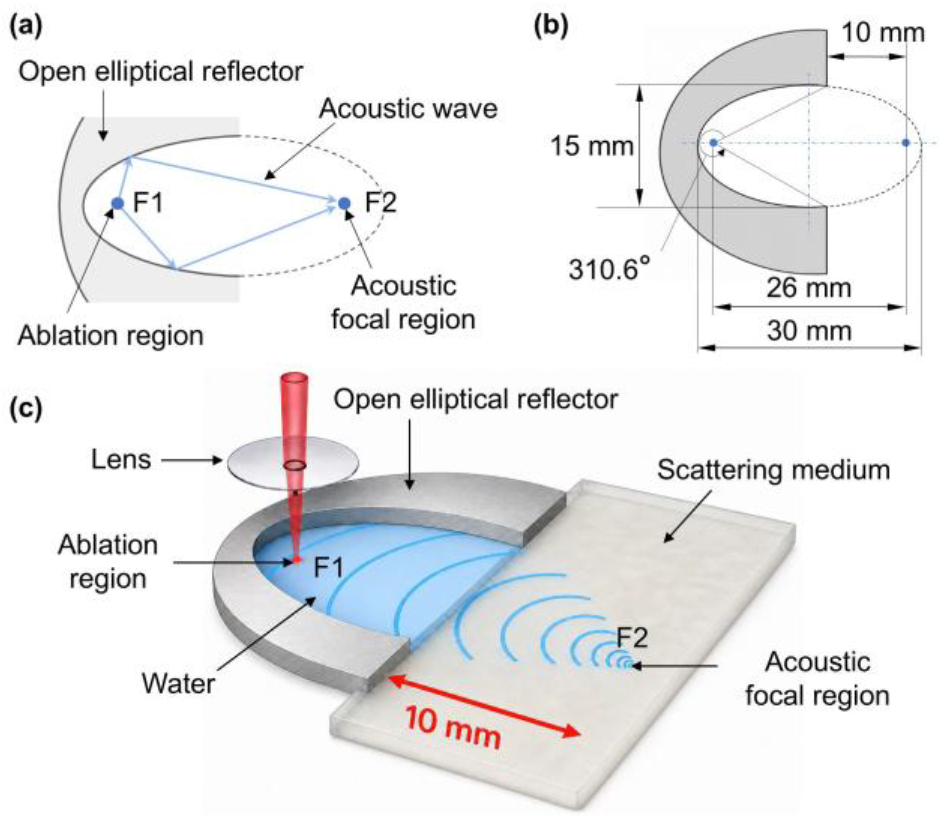
Spatially separated acoustically assisted light guiding using an openellipticalreflector. (a) Acoustic-waveredirection fromtheablation site to the target region. (b) Reflector design. (c) Configuration showing acoustic focusing inside the scattering medium with the ablation site outside.

We fabricated a partially cut open elliptical reflector to redirect and focus laser-induced acoustic waves, as shown in Fig. 1(b). The reflector had a major-axis length of 30 mm, a minor-axis length of 15 mm, and a focal-point separation of 26 (mathematically 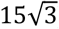) mm. In this design, the reflector opening toward the sample side was set 10 mm from the acoustic focusing position. The opening subtends an angle of 49.4° at the first focal point mathematically. Accordingly, the remaining reflector theoretically covers 310.6° of the full 360° in-plane angular range, corresponding to an ideal geometric coverage of 86.3%. Based on the elliptical geometry, the reflected acoustic path length from the first focal region to the second focal region was designed to be 30 mm, corresponding to an expected arrival time of about 20 µs in water.

In operation, laser ablation at the first focal region generates acoustic waves, which are reflected by the elliptical surface and focused at the second focal region, producing a localized acoustic wave focal region (Fig. 1(c)). Importantly, the ablation region is located outside the sample, whereas the acoustic focal region is generated inside the sample. This geometry allows the acoustic focal region and the associated light transmittance modulation to be controlled by the reflector design, while physically separating the acoustic-wave generation site from the target region.

To verify that the reflector redirected the wave from the separated ablation site and produced localized acoustic focusing on the targeted region, we visualized laser-induced acoustic-wave propagation near the second focal region using partial elliptical aluminum reflector. The reflector and a nonscattering hydrogel target, both 2 mm thick, were sandwiched between two glass windows, and the reflector was filled with water. Fig. 2(a) shows the optical setup. For acoustic wave generation, a pulsed laser with a wavelength of 1064 nm, pulse duration of 5 ns, and pulse energy of 66 mJ (LS-2132UT, LOTIS TII, BY) was focused into the first focal region of the elliptical reflector by an objective lens (10x, NA = 0.25). Acoustic wave propagation around the second focal region was imaged by a time-resolved shadowgraph technique using a 640 nm pulsed laser source (CAVILUX Smart, Cavitar Ltd, FI) with the pulse duration of 20 ns, bandpass filter, and a CMOS camera (CS126MU, Thorlabs, US). The delay time between 1064 nm pulses and 640 nm pulses was changed every 0.01 µs step using a digital delay generator (DG645, Stanford Research Systems, US). Acoustic-wave propagation was isolated by subtracting a background image acquired in the absence of the acoustic wave from each image.

**Fig. 2.**
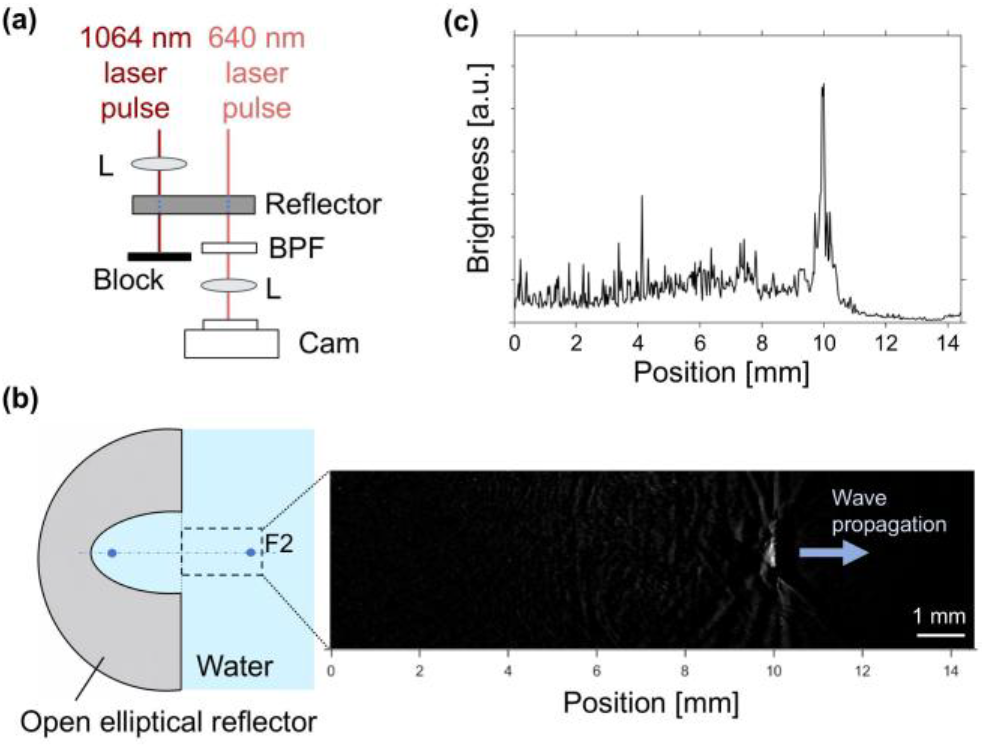
Experimental validation of localized acoustic focusing using an open elliptical reflector. In the side view of the reflector, the focal region F1 is on the left side of the image, and focal region F2 is on the right. L, lens; BPF, band-pass filter; Cam, camera. (a) Shadowgraph setup and reflector geometry. (b) Background-subtracted image showing acoustic-wave propagation near F2. (c) Horizontal brightness profile showing a localized peak 10 mm from the reference edge.

Figure 2(b) shows the acoustic-wave propagation at the delay time 20.03 µs in which the wavefront reached F2, and Fig. 2(c) shows the brightness profile extracted from the rectangular ROI in Fig. 2(b). The wavefront redirected by the open elliptical reflector converges toward the designed acoustic focal region, indicating that the reflector reshapes the acoustic propagation and modulates optical properties at a spatially separated target site.

We next investigated the performance of the proposed method in strongly scattering media and determined the spatial and temporal operating window of acoustically assisted light guiding. The 640-nm pulsed laser was replaced with an 800 nm, 35 fs pulsed laser (ASTRELLA USP-1K-NV, Coherent, Inc., US) because this wavelength is close to the near-infrared excitation wavelength used in multiphoton microscopy. A 2-mm-thick 1% (w/v) intralipid hydrogel was used as a tissue-mimicking scattering phantom, and a 2-mm-thick stainless-steel reflector was used for acoustic wave focusing. The concentration was chosen to reasonably approximate the scattering properties of brain and skin tissues [14,15]. Thus, the selected thickness provided a strongly scattering propagation condition for evaluating the acoustically induced optical modulation. Shadowgraph images were acquired at delays from 19.75 to 20.30 µs after ablation in 0.01 µs steps, with ten repeated measurements at each delay.

Figure 3(a) shows representative images at 19.75, 20.08, and 20.30 µs. The intensity of image initially remained close to the background level. Around 20 µs, a localized optical contrast approximately 20 µm × 230 µm in size appeared near the designed acoustic focal region, indicating localized optical modulation associated with the focused acoustic field.

**Fig. 3.**
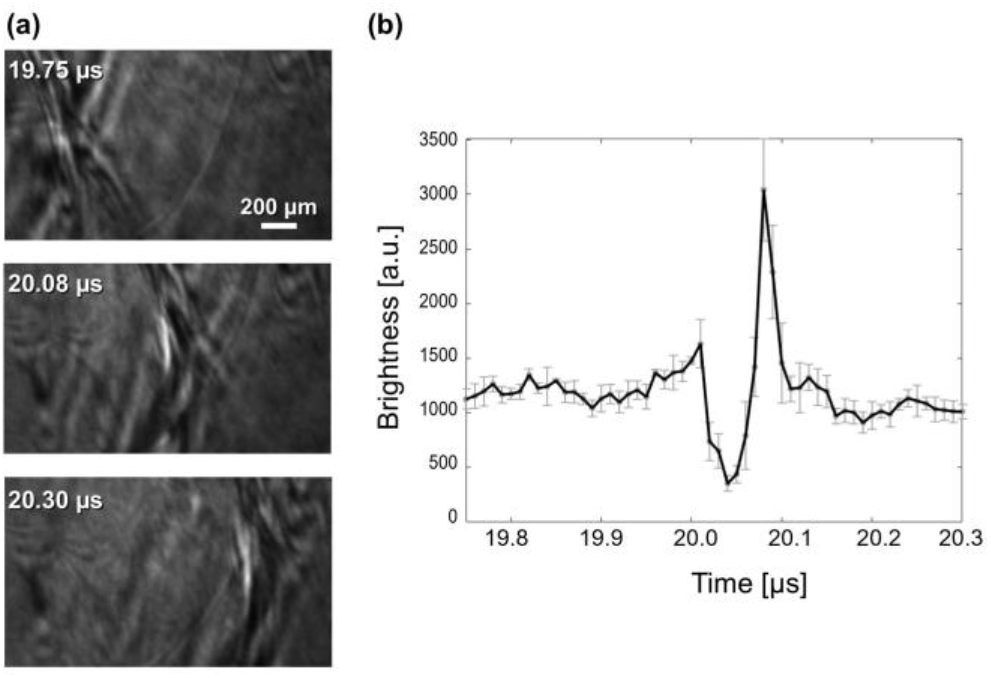
Temporal characterization of acoustic waves-induced optical contrast near the second focal region. (a) Representative images near the second focal region at selected delays after ablation. (b) Temporal ROI brightness profile near the second focal region. The curve and error bars represent the median and median absolute deviation over ten repeated excitations, respectively. The peak appears at 20.08 µs with an FWHM of approximately 18 ns.

The temporal brightness profile extracted from a 10 µm × 100 µm region of interest centered at the designed second focal position is shown in Fig. 3(b). The light transmittance initially decreased upon arrival of the acoustic wave, subsequently increased, and then returned to its initial level. This decrease in light transmittance is attributed to light at the leading edge of the acoustic wavefront being guided toward the pressure peak rather than propagating straight through or scattering radially. This behavior is consistent with a previous study in which acoustic waves were uniformly focused from all directions over 360° [8]. The light transmittance reached its maximum enhancement at 20.08 µs, with a peak brightness 2.57 times the average background intensity. The baseline brightness was determined by averaging the stable portion of the temporal profile. At the position of maximum brightness, the temporal FWHM was calculated relative to this baseline using linear interpolation at the half-maximum crossings, yielding a value of approximately 18 ns. Accordingly, the probe pulse must be synchronized with the arrival of the acoustic wave within this temporal window.

To demonstrate the potential applicability of reflector-based acoustically assisted light guiding to multiphoton microscopy, we performed imaging of two-photon excitation through scattering media. The experimental setup is shown in Fig. 4(a). An 800-nm excitation beam was focused onto the front surface of the hydrogel using a lens with a focal length of 250 mm. Fluorescence emitted from the Rhodamine B layer was collected using an objective lens (10x, NA = 0.45). The collected fluorescence was then relayed by a tube lens (F = 50 mm) and imaged onto a CMOS camera, with a dichroic mirror (638 nm Cut-On) and band-pass filter to reject the 800-nm excitation light. As illustrated in Fig. 4(b), the sample consisted of a 1.5-mm-thick 1% (w/v) intralipid hydrogel and a thin fluorescent layer containing 100 µM Rhodamine B solution behind the hydrogel. A 1.5-mm-thick stainless-steel reflector was used in this configuration. Incident light for two-photon excitation was focused onto the front surface of the hydrogel, then propagated through the scattering layer along the acoustic focal region before reaching the fluorescent layer. By spatially aligning the excitation beam with the acoustic focal region and temporally synchronizing the excitation pulse with the arrival of the focused acoustic wave, we compared fluorescence images acquired with and without synchronized acoustic focusing.

**Fig. 4.**
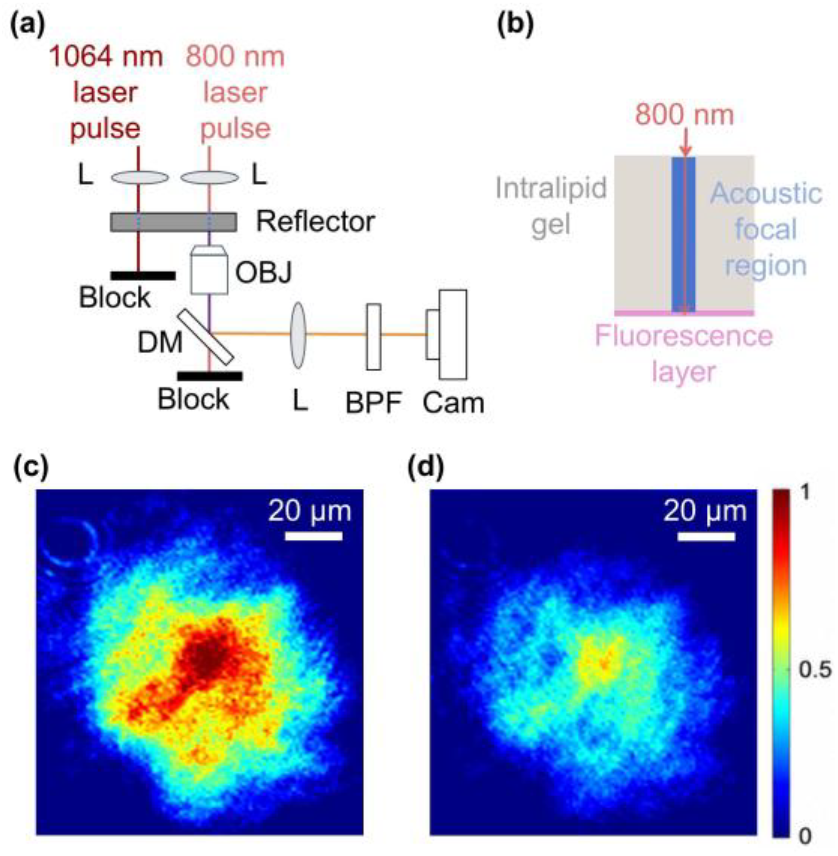
Fluorescence enhancement through a scattering medium using spatially separated laser-induced acoustics. (a) Experimental setup. The focal region F1 is on the left side and the focal region F2 is on the right. A 1064-nm laser pulse generates an acoustic wave at F1, while an 800-nm femtosecond laser excites fluorescence through the acoustic focal region at F2. L, lens; DM, dichroic mirror; OBJ, objective lens; BPF, band-pass filter; Cam, camera. (b) Enlarged view of the excitation path through the 1.5-mm-thick intralipid hydrogel to the fluorescent layer. (c, d) Fluorescence images with and without synchronized acoustic-wave-assisted light guiding, respectively. Both images are displayed using a common intensity scale normalized to the synchronized image.

As shown in Fig. 4(c) and Fig. 4(d), the fluorescence signal acquired without synchronized acoustic focusing was weaker because the 800-nm excitation light was scattered while propagating through the intralipid hydrogel. In contrast, when the excitation pulse was synchronized with the acoustic-focusing window, the fluorescence intensity increased and the bright region became more pronounced. This enhancement suggests that the acoustically induced light transmittance modulation near the second focal region improved the delivery of excitation light to the 100 μM Rhodamine B layer. These results demonstrate that the acoustic focal region can be spatially aligned with the excitation beam and that the optical pulse can be temporally synchronized with the transient light transmittance modulation, thereby enhancing fluorescence excitation through the scattering medium without moving the sample.

In conclusion, we demonstrated an acoustically assisted light-guiding method based on remotely generated acoustic waves and an open elliptical reflector. Laser-induced acoustic waves generated outside the sample were redirected by the reflector and focused at a spatially separated second focal region, enabling formation of a localized acoustic focal region without moving the sample itself. The reflected acoustic waves produced localized and transient modulation of light transmission at the designed target position. Synchronizing optical excitation with this modulation enhanced two-photon excitation through a scattering medium, demonstrating the potential of this approach for spatially controlled light delivery in scattering media.

Future work on adapting this reflector-based acoustic-focusing approach to biological imaging will need to address two main challenges. First, tissue heterogeneity may distort the propagating acoustic field and degrade focusing at the target. Because the acoustic waves are generated optically, such distortions could potentially be compensated by tailoring the laser profile and thereby modifying the spatial characteristics of the acoustic source. Second, spatial scanning of the optical target would require not only conventional scanning of the excitation beam but also repositioning of the acoustic focus. This may be achieved, within a limited range, by shifting the laser-induced acoustic source away from the exact first focal position, thereby displacing the corresponding acoustic focus. The feasibility, correction capability, and scanning range of these approaches will need to be evaluated through simulations and experiments. If realized, such control may facilitate localized light delivery in scattering tissue for applications including multiphoton fluorescence imaging, optogenetic stimulation [16,17], and photodynamic therapy [18].

## Data availability

Data underlying the results presented in this paper are not publicly available at this time but may be obtained from the authors upon reasonable request.

## References

1. F. Helmchen and W. Denk, “Deep tissue two-photon microscopy,” Nat. Methods 2, 932–940 (2005).

2. R. P. J. Barretto, T. H. Ko, J. C. Jung, et al., “Time-lapse imaging of disease progression in deep brain areas using fluorescence microendoscopy,” Nat. Med. 17, 223–228 (2011).

3. G. Cui, S. B. Jun, X. Jin, et al., “Deep brain optical measurements of cell type–specific neural activity in behaving mice,” Nat. Protoc. 9, 1213–1228 (2014).

4. Q. Guo, J. Zhou, Q. Feng, et al., “Multi-channel fiber photometry for population neuronal activity recording,” Biomed. Opt. Express 6, 3919–3931 (2015).

5. R. Horstmeyer, H. Ruan, and C. Yang, “Guidestar-assisted wavefront-shaping methods for focusing light into biological tissue,” Nat. Photonics 9, 563–571 (2015).

6. I. M. Vellekoop and A. P. Mosk, “Focusing coherent light through opaque strongly scattering media,” Opt. Lett. 32, 2309–2311 (2007).

7. M. Chamanzar, M. G. Scopelliti, J. Bloch, et al., “Ultrasonic sculpting of virtual optical waveguides in tissue,” Nat. Commun. 10, 92 (2019).

8. A. Ishijima, U. Yagyu, K. Kitamura, et al., “Nonlinear photoacoustic waves for light guiding to deep tissue sites,” Opt. Lett. 44, 3006–3009 (2019).

9. M. N. Cherkashin, C. Brenner, G. Schmitz, et al., “Transversally travelling ultrasound for light guiding deep into scattering media,” Commun. Phys. 3, 180 (2020).

10. P. Ricci, M. Colom, B. Mestre-Torà, et al., “Photoacoustics for direct light-guiding inside transparent and scattering media,” Laser Photonics Rev. 19, 2401122 (2025).

11. M. Taniguchi, H. Fukuoka, T. Nitta, et al., “Numerical calculation to elucidate effect of open elliptical reflector shape on underwater shock wave focusing phenomena,” Jpn. J. Appl. Phys. 64, 042003 (2025).

12. Y. Zhou and P. Zhong, “The effect of reflector geometry on the acoustic field and bubble dynamics produced by an electrohydraulic shock wave lithotripter,” J. Acoust. Soc. Am. 119, 3625–3636 (2006).

13. J. I. Iloreta, Y. Zhou, G. N. Sankin, et al., “Assessment of shock wave lithotripters via cavitation potential,” Phys. Fluids 19, 086103 (2007).

14. T. Vo-Dinh, ed., Biomedical Photonics Handbook (CRC Press, 2003).

15. P. Lai, X. Xu, and L. V. Wang, “Dependence of optical scattering from Intralipid in gelatin-gel based tissue-mimicking phantoms,” J. Biomed. Opt. 19, 035002 (2014).

16. A. M. Packer, B. Roska, and M. Häusser, “Targeting neurons and photons for optogenetics,” Nat. Neurosci. 16, 805–815 (2013).

17. Y. Shin, M. Yoo, H.-S. Kim, et al., “Characterization of fiber-optic light delivery and light-induced temperature changes in a rodent brain for precise optogenetic neuromodulation,” Biomed. Opt. Express 7, 4450–4471 (2016).

18. M. M. Kim and A. Darafsheh, “Light sources and dosimetry techniques for photodynamic therapy,” Photochem. Photobiol. 96, 280–294 (2020).

